# Gini coefficients for measuring the distribution of sexually transmitted infections among individuals with different levels of sexual activity

**DOI:** 10.1101/438127

**Authors:** Sandro Gsteiger, Nicola Low, Pam Sonnenberg, Catherine H Mercer, Christian L Althaus

## Abstract

**Objectives:** Gini coefficients have been used to describe the distribution of *Chlamydia trachomatis* (CT) infections among individuals with different levels of sexual activity. The objectives of this study were to investigate Gini coefficients for different sexually transmitted infections (STIs), and to determine how STI control interventions might affect the Gini coefficient over time.

**Methods:** We used population-based data for sexually experienced women from two British National Surveys of Sexual Attitudes and Lifestyles (Natsal-2: 1999-2001; Natsal-3: 2010-2012) to calculate Gini coefficients for CT, *Mycoplasma genitalium* (MG), and human papillomavirus (HPV) types 6, 11, 16 and 18. We applied bootstrap methods to assess uncertainty and to compare Gini coefficients for different STIs. We then used a mathematical model of STI transmission to study how control interventions affect Gini coefficients.

**Results:** Gini coefficients for CT and MG were 0.33 (95% confidence interval (CI): 0.18-0.49) and 0.16 (95% CI: 0.02-0.36), respectively. The relatively small coefficient for MG suggests a longer infectious duration compared with CT. The coefficients for HPV types 6, 11, 16 and 18 ranged from 0.15-0.38. During the decade between Natsal-2 and Natsal-3, the Gini coefficient for CT did not change. The transmission model shows that higher STI treatment rates are expected to reduce prevalence and increase the Gini coefficient of STIs. In contrast, increased condom use reduces STI prevalence but does not affect the Gini coefficient.

**Conclusions:** Gini coefficients for STIs can help us to understand the distribution of STIs in the population, according to level of sexual activity, and could be used to inform STI prevention and treatment strategies.

**Key messages:**

- The Gini coefficient can be used to describe the distribution of STIs in a population, according to different levels of sexual activity.
- Gini coefficients for *Chlamydia trachomatis* (CT) and human papillomavirus (HPV) type 18 appear to be higher than for *Mycoplasma genitalium* and HPV 6, 11 and 16.
- Mathematical modelling suggests that CT screening interventions should reduce prevalence and increase the Gini coefficient, whilst condom use reduces prevalence without affecting the Gini coefficient.
- Changes in Gini coefficients over time could be used to assess the impact of STI prevention and treatment strategies.

## INTRODUCTION

Understanding how sexually transmitted infections (STIs) are distributed among individuals is important both from a biological and from a public health perspective. Differences in STI prevalence within a population, between groups with varying levels of sexual activity, can provide information about biological and epidemiological characteristics of the infection. For example, an STI with a long infectious duration, such as human papillomavirus (HPV), will tend to be spread more evenly across a population than an STI with a short infectious duration, such as Neisseria gonorrhoeae (NG). NG is more concentrated in a small subgroup of individuals with high sexual activity. Such ideas were initiated in the late 1970s and led to the concept of the ‘core group’ [1]. In 1990, Brunham & Plummer [2] inferred the size of core groups for various STIs from the biological parameters that describe transmissibility and infectious duration, and discussed the implications for selecting adequate STI control strategies.

The Gini coefficient can be used to quantify the degree of concentration of an STI in a population. Originally introduced for describing inequalities in income distributions [3], the Gini coefficient provides a general tool to measure the distribution of a disease outcome in relation to an exposure variable or risk factor [4], such as the geographic location or sexual behaviour. A Gini coefficient of zero denotes perfect equality where an infection is equally distributed across a population. For infections that are concentrated in specific subpopulations, the Gini coefficient can increase up to a maximal value of one. The Lorenz curve is a visual representation of the cumulative distribution of a disease when ordered according to the risk factor [5]. The diagonal line on a Lorenz curve plot denotes perfect equality, e.g., every subpopulation has the same prevalence of an STI.

Several groups have used Gini coefficients and Lorenz curves to describe how *Chlamydia trachomatis* (CT), NG, syphilis or herpes are distributed across different geographical regions in Canada [6], the UK [7] and the US [8–10]. These findings have helped to summarise inequalities in STI distributions, assess the suitability of geographically targeted interventions, and provide insights into the epidemic phases of the STIs over time. Chen et al. [11] were the first to apply the concept of Gini coefficients in mathematical transmission models. They proposed a metapopulation modelling framework that better captures the sociogeographic epidemiology of NG and compared the resulting Gini coefficients with empirical estimates.

The way in which modifiable factors, such as sexual activity and STI control interventions affect the Gini coefficient has been less-well studied. Previously, we described the distribution of CT infections among individuals with different sexual activity using Lorenz curves and Gini coefficients to calibrate dynamic transmission models [12]. We used data from the second British Survey of Sexual Attitudes and Lifestyles (Natsal-2, 1999-2001) [13], which included CT test results (samples were later tested for HPV [14]). The most recent survey (Natsal-3, 2010-2012) also provides information on HPV and Mycoplasma genitalium (MG) positivity and offers a unique opportunity to study relationships between sexual behaviour and STI prevalence [15–17].

This study had two objectives. First, we wanted to estimate and compare the Gini coefficients for different STIs and over time. Second, we wanted to obtain general insights as to what extent changes in Gini coefficients over time can provide information about the effects of control interventions for STIs, using a mathematical transmission model.

## METHODS

### Data

Natsal-3 is a population-based probability sample survey of sexual attitudes and lifestyles conducted in Britain (England, Scotland, and Wales) and carried out from 2010 to 2012 [18,19]. The sample consists of 15162 women and men aged 16-74 years. A subsample of participants aged 16-44 years who reported at least one sexual partner over their lifetime were asked to provide urine samples, resulting in laboratory confirmed STI test results from 2665 women and 1885 men [15]. Urine was tested for the presence of CT, MG, type-specific HPV, NG, and HIV antibodies. To compare Gini coefficients for CT over time, we also used data from the Natsal-2 survey, which was carried out in 1999-2001 [20]. This survey includes 11,161 women and men aged 16-44 years. Urine samples for ligase chain reaction (LCR) testing for CT were available for a subset of 2055 and 1474 sexually active women and men aged 18-44 years [13]. Unless otherwise stated, we used the respective subpopulations that provided urine samples for our analysis from both surveys. We applied individual weights to all variables to adjust for unequal selection probabilities and to correct for the age, gender and regional profiles in the survey sample. For simplicity, we did not include same-sex contacts for the sexual behaviour variables because only 2.2% of the population reported a new same-sex partner during time period covered by Natsal-3. The full datasets of both surveys are available from the UK Data Archive at the University of Essex (http://data-archive.ac.uk).

### Statistical analysis

We used Lorenz curves to plot the cumulative proportion of specific STI infections (*y*_*i*_) as a function of the cumulative proportion of the population (*x*_*i*_), after population sub-groups *i* (*i* = 1,2, …, *n*) have been ranked according to their level of sexual activity. The Gini coefficient is defined as the area between the line of equality and the Lorenz curve over the total area below the line of equality:

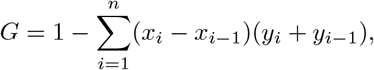

where *x*_0_ = *y*_0_ = 0 and *x*_*n*_ = *y*_*n*_ = 1.

We derived Lorenz curves and estimated Gini coefficients for CT, MG, and HPV types 6, 11, 16 and 18. We focused on these four HPV genotypes because they are present in the widely used quadrivalent vaccine and are frequently considered in dynamic transmission models [21]. We did not include NG and HIV in our analysis because of the small numbers of positive tests. We used the number of new opposite-sex partners in the last year as the exposure variable summarising sexual activity because of its strong association with STI prevalence [12], and frequent use to parameterise dynamic transmission models [22]. Owing to the larger sample size for female respondents and a potential bias resulting from lower test sensitivity to detect HPV infections in male urine [15], we focused our analysis on women and provide a separate analysis for men in the Supplementary material. We constructed bootstrap confidence intervals (CIs) for the Gini coefficients and point-wise bootstrap confidence bands for the Lorenz curves [8–10,23]. We calculated the 2.5th and 97.5th percentiles from 1000 bootstrapped Gini coefficients and Lorenz curves.

### Transmission model

We adapted a previously described mathematical model of STI transmission [24] to investigate how changes in infectious duration and transmissibility affect the prevalence and the Gini coefficient of an STI in a simulated population. We stratified the population according to sexual activity [1,25], i.e., we assumed *n* different sexual activity classes with 0,1,2,…, *n*-1 new opposite-sex partners per year. For simplicity, we assumed that sexual activity and the natural history and transmission of the infection are the same in women and men. The SIS (susceptible-infected-susceptible) transmission dynamics can be described by the following ordinary differential equation (ODE):

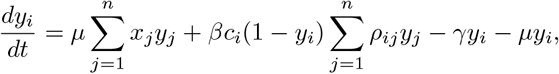

where *y*_*i*_ is the proportion of individuals in sexual activity class *i* who are infected, and *x*_*i*_ is the proportion of all individuals who belong to sexual activity class *i*. The first and last term of the equation describe how individuals can change their sexual activity class at rate *μ* and be redistributed to either the same or another sexual activity class proportional to the size of sexual activity classes [24,26]. The middle terms describe the process of transmission and the clearance of infection at rate *γ*. Susceptible individuals 1 − *y*_*i*_ have an average of *c_i_* new opposite-sex partners per year. *β* is the per partnership transmission probability and *y_j_* is the probability that a partner in sexual activity class *j* is infected. *ρ*_*ij*_ represents the elements of the sexual mixing matrix [27]

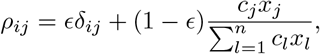

where *δ*_*ij*_ denotes the Kronecker delta (it is equal to 1 if *i* = *j* and to 0 otherwise).

The proportion of individuals in sexual activity classes *x*_*i*_ with *c*_*i*_ ∈ 0, 1, 2, …, *n* − 1 new opposite-sex partners per year were based on Natsal-2. All data for 18-44 year old women and men were pooled and weighted. Since the transmission model is primarily used for illustrative purposes, we did not include changes in sexual behaviour, between Natsal-2 and Natsal-3, which were small [19]. We set *μ* = 1 per year as described previously [24,26], and set *ϵ* = 0, i.e., we assumed random proportional mixing between different sexual activity classes [27]. We then chose a particular combination of the infectious duration (1/*γ*) and the per partnership transmission probability (*β*) and ran the model into endemic equilibrium by numerically integrating the ODEs in the R software environment for statistical computing [28] using the function *ode* from the package *deSolve* We calculated the prevalence and Gini coefficient as described above. We repeated this process for various combinations of the infectious duration and the per partnership transmission probability, which allowed us to map between the (unobservable) model parameters and the (observable) summary measures prevalence and Gini coefficient.

We used the model to investigate two CT control interventions. First, we assumed that an increase in screening coverage, which aims to detect asymptomatic CT cases and limits transmission by early treatment, will reduce the overall infectious duration by 10%. Second, we modelled the effects of an educational campaign that leads to a behaviour change and an increase in condom use by reducing the per partnership transmission probability by 10%.

## RESULTS

### Lorenz curves and Gini coefficients

The Lorenz curves for MG and HPV types 6, 11 and 16 are closer to the diagonal line than the Lorenz curves for CT and HPV 18 (figure 1A). This indicates that CT and HPV-18 in women are more strongly associated with the number of new opposite-sex partners in the last year than MG and the other type-specific HPV. The Gini coefficients mirror this observation and are higher for CT (0.33) and HPV 18 (0.38) than for MG and the different HPV types (≤ 0.22) (table 1). However, the bootstrapped confidence intervals for the Lorenz curves are wide (figure 1B), resulting in considerable uncertainty in the estimated Gini coefficients. The Lorenz curves for CT for the two survey periods of Natsal-2 (1999-2001) and Natsal-3 (2010-2012) are similar (figure 1C) with a Gini coefficient in Natsal-2 of 0.41 (95% CI: 0.18-0.61) and in Natsal-3 of 0.33 (95% CI: 0.18-0.49).

**Figure 1.**
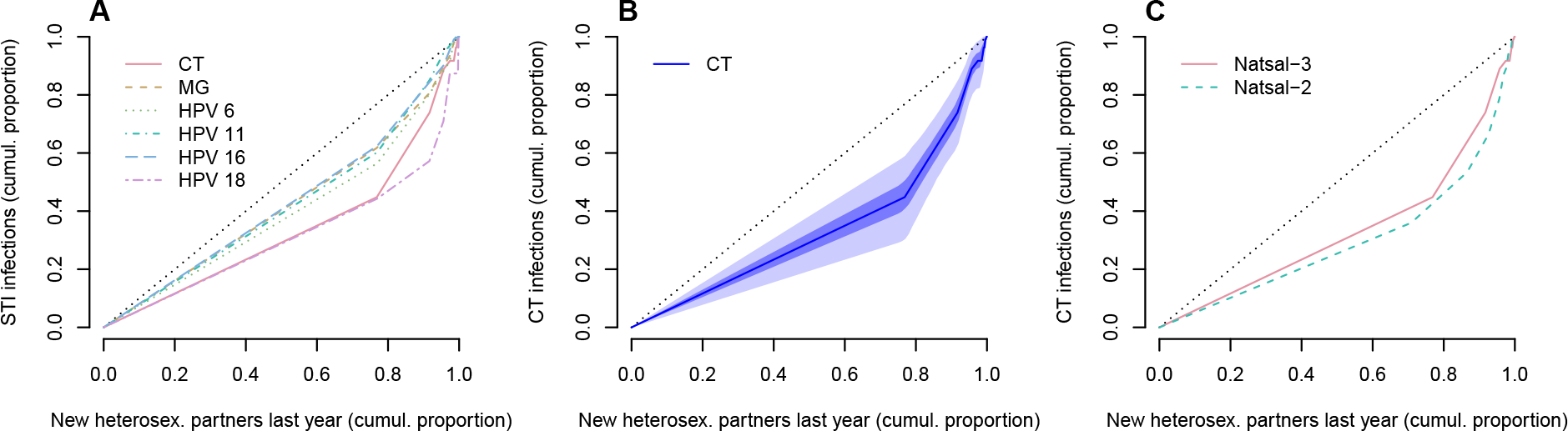
Lorenz curves representing the cumulative proportion of STI infections in women as a function of the cumulative proportion of the population, after population sub-groups have been ranked by the number of new opposite-sex partners in the last year. (A) Lorenz curves for different STIs. Data: Natsal-3. (B) Uncertainty around Lorenz curve for CT. The grey areas represent point-wise 50% (dark grey) and 95% (light grey) confidence bands. Data: Natsal-3. (C) Comparison of Lorenz curves for CT between Natsal-2 (dashed line) and Natsal-3 (solid line). In all graphs, the diagonal line (black dotted line) denotes perfect equality, i.e., an equal dispersion of the infection across population sub-groups.

**Table 1.** Estimated Gini coefficients for different sexually transmitted infections in women

| Infection | Gini coefficient | 95% confidence interval (CI) |
| --- | --- | --- |
| Chlamydia trachomatis | 0.33 | 0.18 - 0.49 |
| Mycoplasma genitalium | 0.16 | 0.02 - 0.36 |
| HPV 6 | 0.22 | 0.02 - 0.43 |
| HPV 11 | 0.17 | 0.04 - 0.31 |
| HPV 16 | 0.15 | 0.04 - 0.27 |
| HPV 18 | 0.38 | 0.10 - 0.64 |

The results from the transmission model show an intricate relationship between the Gini coefficient, infection prevalence, infectious duration and per partnership transmissibility (figure 2A, dashed grid). STIs with a short infectious duration (e.g., 1 year) are primarily transmitted between individuals with a high number of opposite-sex partners and are characterised by a high Gini coefficient. Longer infectious durations increase prevalence, facilitate STI transmission between individuals with a low number of opposite-sex partners, and consequently decrease the Gini coefficient. Interestingly, different values for the per partnership transmission probability only influence prevalence but do not affect the Gini coefficient. These insights from the transmission model potentially allow the inference of biological parameters for the different STIs in Natsal-3 (figure 2A, coloured dots). Although the confidence intervals and the associated uncertainty are large, CT and HPV-18 seem to be consistent with an infectious duration between 1 and 2 years. MG and the other HPV types are consistent with longer infectious durations.

**Figure 2.**
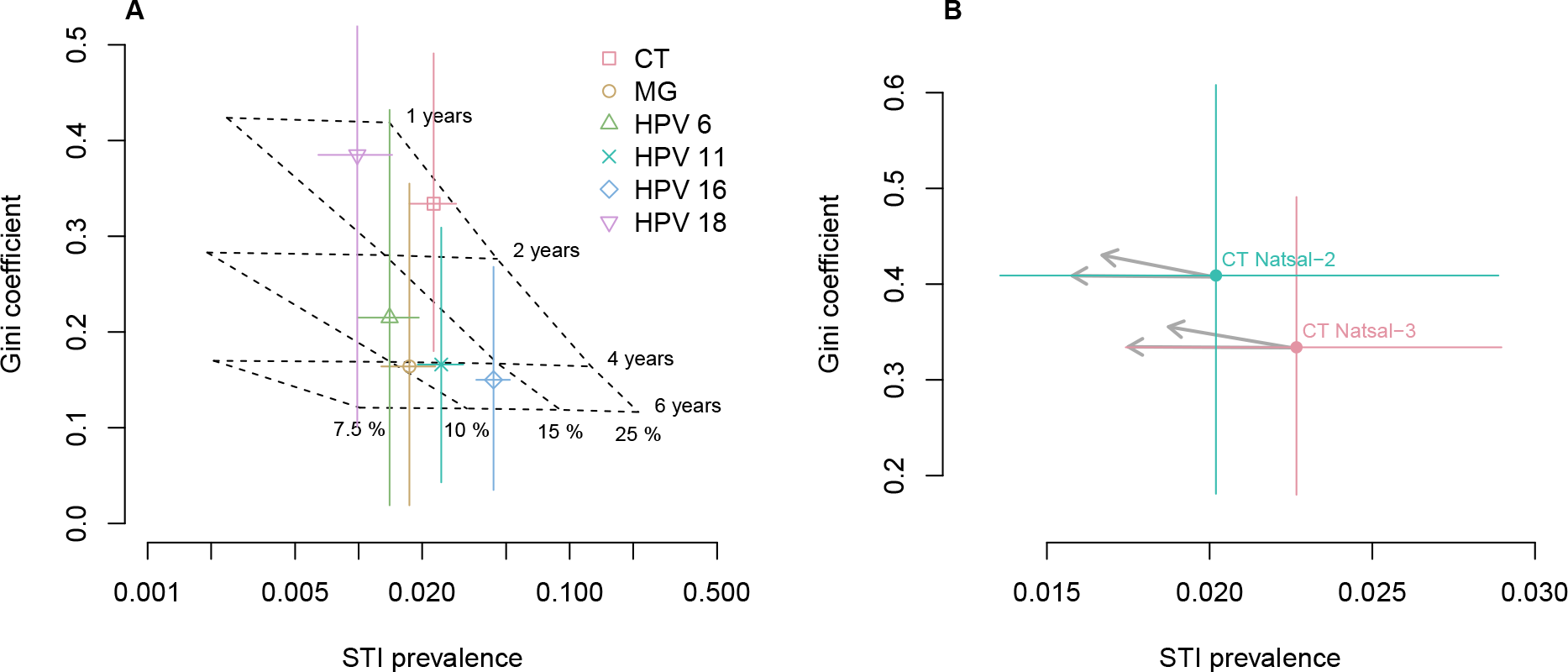
Relationship between Gini coefficient, STI prevalence, infectious duration and transmissibility. (A) Gini coefficients and STI prevalence for women in Natsal-3 (coloured dots). Modelled values for different combinations of the infectious duration and the per partnership transmission probability are projected on the graph (dashed grid). (B) Expected impact of control measures on Gini coefficients and prevalence of female CT in Natsal-2 and Natsal-3. The black arrows denote a 10% reduction in the per partnership transmission probability (horizontal arrows) or the infectious duration (diagonal arrows).

### STI control interventions

We used the transmission model to examine how two different control interventions for CT affect the Gini coefficient and infection prevalence. First, we assumed that the increase in CT screening between the two survey periods of Natsal-2 (1999-2001) and Natsal-3 (2010-2012) through the National Chlamydia Screening Programme (NCSP) [29] has resulted in a reduction in the overall infectious duration. The transmission model suggests a concurrent drop in prevalence with a higher Gini coefficient (figure 2B). The point estimates for the prevalence and Gini coefficient in Natsal-2 and Natsal-3 move in the opposite direction. The expected changes of a 10% reduction in the infectious duration would be relatively small, however, and may remain within the 95% CIs of the point estimates. Second, we assumed that an increase in condom use results in a 10% reduction in the per partnership transmission probability. Such an intervention would then be expected to reduce the prevalence of CT but would not affect the Gini coefficient.

## DISCUSSION

Building upon earlier work [12], we constructed Lorenz curves and estimated Gini coefficients in women to investigate how different STIs are distributed according to sexual activity. Gini coefficients for CT and HPV 18 appear to be higher than for MG and HPV 6, 11 and 16. We found no evidence that the Gini coefficient for CT changed between the two survey periods of Natsal-2 and Natsal-3. Using a mathematical model of STI transmission, we found that a CT screening intervention should reduce prevalence and increase the Gini coefficient, whilst condom use reduced prevalence but did not affect the Gini coefficient.

A main strength of this study was the availability of Natsal-2 and Natsal-3, two very large data sets that measure both STI positivity and self-reported sexual behaviour in probability samples of the British general population. These comprehensive data sets allow comparison between different STIs and over time. Calculating infection prevalence and Gini coefficients is straightforward if suitable data are available. In contrast, obtaining biological parameters that determine the transmission dynamics, such as the infectious duration or the transmission probability, are notoriously difficult to obtain [30,31]. In this study, we used a mathematical model of STI transmission with a detailed description of different sexual activity classes [24] to explore the relationship between Gini coefficient, infection prevalence, infectious duration and transmissibility.

There are several limitations to this study. First, despite the large overall sizes of the two Natsal surveys, the relatively low prevalence of STIs in the general population [13–16] resulted in relatively large uncertainties in the Lorenz curves and the corresponding Gini coefficients, particularly for males (see Supplementary material). Owing to the small sample sizes, we pooled data over all age groups and restricted our analyses to the whole survey population. Investigating sex-or age-specific differences in Gini coefficients would certainly be interesting but is currently not feasible. Second, the limited sample size did not allow us to calculate Gini coefficients for NG and HIV, which have a low prevalence, and arguably high Gini coefficient [15]. Third, the comparison of prevalence and Gini coefficients of CT between Natsal-2 and Natsal-3 should also be treated with caution. The two surveys used different laboratory tests and were not powered to detect changes in CT prevalence [15]. Fourth, we defined the exposure variable as the number of new opposite-sex partners in the last year. Using other exposure variables for our analysis would obviously affect the Lorenz curves and the Gini coefficients, but would not necessarily be applicable for the mapping between model parameters and the summary measures. In addition, mathematical modelling necessarily involves several assumptions and simplifications. As in the data analysis, we did not stratify the population according to age, assumed the sexual behaviour in women and men to be the same, and considered the general population of those reporting sex with opposite-sex partners as a whole. Changes in sexual behaviour between Natsal-2 and Natsal-3 were minimal [19], so we did not take these into account. We assumed fully proportional mixing which typically results in the best description of infection prevalence in different sexual activity classes in models with a constant per partnership transmission probability [22,24,27]. We also did not consider sex-specific differences in the infectious duration and the transmissibility of the various STIs, which might limit the application of Gini coefficients for the inference of these parameters. Finally, we assumed that individuals who clear an infection can become reinfected, although CT and type-specific HPV infections might confer temporal immunity [32,33].

The calculated Gini coefficients and prevalence of CT and HPV 18 in women in Natsal-3 suggest an infectious duration of 1-2 years, which is in good agreement with previous estimates [30,34–36]. Our mapping indicates that the infectious durations for the other high-risk HPV type 16 and low-risk HPV types 6 and 11 could be longer than 2 years. This interpretation would contrast with previous studies that estimate similar infectious durations for HPV-16 and HPV-18 [34,35], and shorter infectious durations of less than a year for HPV 6 and 11 [34]. The discrepancy could be a result of ignoring the effects of temporal immunity to reinfection. The infectious duration for the other bacterial STI, MG, seems to be longer than for CT. There is considerable uncertainty regarding the infectious duration of MG. One analysis, which used data from a study of female students in London (UK), estimated the mean infectious duration at 15 months [37], which is in the same range as CT. The per partnership transmission probability is a highly model-dependent parameter and depends on the type of sexual partnerships that are considered. It is maybe not surprising that our mapping, which suggests relatively low per partnership transmission probabilities of 10% to 25%, is not consistent with estimates from other modelling studies [31,33].

Our findings allowed us to interpret differences in Gini coefficients and changes over time for different STIs, illustrating that Gini coefficients can serve beyond their original role as simple statistical measures of exposure-outcome associations. We found that changes in the transmission probability only influence infection prevalence. This means that while decreasing the transmission probability (e.g., through increased condom use) decreases the overall burden of an STI, it does not affect how an STI is distributed among individuals with different sexual activity. Hence, the target groups for future control interventions should remain the same. In contrast, we showed that changes in the infectious duration (e.g., through increased testing and treatment uptake) affects both the prevalence and Gini coefficient of an STI. A stronger concentration of the infections among individuals with increased sexual activity would require a change in the target groups for control interventions.

In summary, our study illustrates that the Gini coefficient for measuring the distribution of an STI among individuals with different sexual activity represents a simple proxy measure, which combines epidemiological and behavioural data. We suggest that estimating Gini coefficients, in combination with mathematical modelling, has the potential to make inference of biological parameters that determine STI transmission and to assess the impact of control measures.

## Acknowledgements

Natsal-3 is a collaboration between University College London, London School of Hygiene and Tropical Medicine, National Centre for Social Research, Public Health England, and the University of Manchester. We thank the study participants, the team of interviewers from NatCen Social Research, and operations and computing staff from NatCen Social Research. The Natsal study was approved by the Oxfordshire Research Ethics Committee A (Ref: 10/H0604/27).

## Contributors

SG, NL and CLA designed the study and wrote the manuscript. PS and CHM provided data. SG and CLA analysed the data. All authors contributed to the interpretation of the results, commented on the manuscript, and approved the final version of the manuscript.

## Funding

Natsal-3 was supported by grants from the Medical Research Council (G0701757); and the Wellcome Trust (084840); with contributions from the Economic and Social Research Council and Department of Health. SG and CLA were funded by the Swiss National Science Foundation (grants 135654 and 136737, respectively).

## Competing interests

The authors declare no competing interests.

